# Color tuning of neurons in face patches of macaque inferior temporal cortex

**DOI:** 10.1101/863779

**Authors:** Marianne Duyck, Tessa J. Gruen, Lawrence Y. Tello, Serena Eastman, Joshua Fuller-Deets, Bevil R. Conway

## Abstract

Previous work has shown that under viewing conditions that break retinal mechanisms for color, one class of objects appears paradoxically colored: faces, and they look green. Interpreted within a Bayesian-observer framework, this observation makes the surprising prediction that face-selective neurons are sensitive to color and weakly biased for colors that elicit L>M cone activity (warm colors). We tested this hypothesis by measuring color-tuning responses of face-selective cells in alert macaque monkey, using fMRI-guided microelectrode recording of the middle and anterior face patches and carefully color-calibrated stimuli. The population of face-selective neurons showed significant color tuning when assessed using images that preserved the luminance contrast relationships of the original face photographs. A Fourier analysis of the color-tuning responses uncovered two components. The first harmonic was biased towards the L>M colors, consistent with the prediction. Interestingly, the second harmonic aligned with the S-cone cardinal axis, which may relate to the computation of animacy by IT cells.

**Significance:** The results provide the first quantitative measurements of the color tuning properties of face-selective neurons. The results provide insight into the neural mechanisms that could support the role of color in face perception.

---

Color is neither necessary nor sufficient for face recognition: People can identify faces in black and white photographs and are impaired at recognizing faces in pictures that have color contrast but lack luminance contrast (Sinha et al., 2006). These observations suggest that the neural machinery for face recognition is color blind (Webster and MacLeod, 2011), an idea that is supported by adaptation experiments, which fail to show interactions between face and color processing (Yamashita et al., 2005). Because color does not appear to contribute to face recognition, most behavioral and neuronal studies of face perception have used achromatic stimuli (Kanwisher et al., 1997; Ohayon et al., 2012). Exhaustive work has uncovered in detail the role of luminance contrast in generating the face selectivity observed in face cells (Ohayon et al., 2012).

The work discussed above might be taken to imply that color plays no role in face perception. Despite the lack of neurophysiological evidence, there is behavioral evidence that color and face processing is linked in one specific way. Under monochromatic viewing conditions, which disable retinal mechanisms for color, faces, and only faces, have an odd color: they appear green (Hasantash et al., 2019). The chromatic signals that give rise to this face-specific paradoxical color perception are selective for the L-M, so-called red-green, chromatic direction. Thus the paradoxical percept is not simply complementary to the normal colors of faces, which vary widely (reflecting variable contributions of melanin and hemoglobin), but restricted to one chromatic direction. The behavioral results in Hasantash et al (2019) were observed when people looked at either African American or Caucasian faces, suggesting that red-green face-color signals are seen regardless of race. One speculation is that the behavioral results reflect the relative importance of dynamic facial color signals, which are predominantly driven by hemoglobin. Regardless, the paradoxical face-color interaction makes an unexpected prediction about the color responses of face selective neurons: that the population is not only sensitive to chromatic information, but also weakly biased towards the L pole in color space. This hypothesis emerges from the best account of similarly “anti-Bayesian” phenomena: Wei and Stocker showed that these phenomena can be understood in the context of a Bayesian observer model constrained by efficient coding (Wei and Stocker, 2015). Their model makes a prediction that the neural population underlying the perceptual phenomena shows a weak bias reflecting the prior. In the cases of faces, the behavioral work suggests that the prior is not simply for natural versus unnatural colors of faces, but specifically for the L-M chromatic axis. The distinction is important because the natural colors of faces vary among people and monkeys, and any given face will elicit activation of all three cone types. Because a likely candidate for encoding face information are face-selective neurons of inferior temporal cortex, we hypothesized that these neurons encode this L>M prior. Is there any evidence for this hypothesis in the literature?

Many cells in macaque temporal cortex can show both complex shape selectivity and color sensitivity (Edwards et al., 2003; Komatsu and Ideura, 1993). To the extent it has been examined, measurements of color responses in face cells have been inconclusive. Of the two studies, one found no sensitivity to color among face cells (Perrett et al., 1982). The other seminal paper found that of 22 face-selective cells in IT, some had slightly higher firing rates for naturally colored face photographs compared to un-naturally colored face photographs. The results in Edwards et al provided the first clues that face cells are responsive to color. But those data are insufficient to test the hypothesis for two reasons. First, responses were only measured to two colors, precluding an evaluation of the color-tuning function of the neurons. And second, it is not known where in IT relative to functional landmarks the face cells studied by Edwards were located. This is an important issue because it is now known that many face-selective cells are clustered within face patches (Tsao et al., 2006), segregated from color-biased domains, in humans and macaque monkeys (Conway, 2018; Lafer-Sousa and Conway, 2013; Lafer-Sousa et al., 2016). Face-selective neurons are clustered in face patches but also found outside face patches (Aparicio et al., 2016). It is possible that the cells of the conflicting studies (Edwards et al., 2003; Perrett et al., 1982) were located in different functionally defined domains. No study has measured color responses face cells within the face patches. Indeed no study has measured color tuning systematically in face-selective neurons in any part of IT. Confronted by strong evidence that the selectivity of face cells is driven entirely by luminance contrast (Ohayon et al., 2012), the possibility that face cells could encode specific color signals has been overlooked. And yet the gross lack of information about the color properties of face cells is surprising given the potential importance of face color to social interactions. Here we addressed this fill this gap in knowledge by testing the specific hypothesis presented by the behavioral results in Hasantash et al. Do face-selective cells show a bias for L>M colors?

## Results

Functional magnetic resonance imaging was used to identify regions of inferior temporal cortex that were more responsive to faces than to bodies and fruits, a standard contrast used to identify face patches (Lafer-Sousa and Conway, 2013; Tsao et al., 2006). We targeted microelectrodes to the ML face patch in two monkeys (M1 and M2) and the AL face patch in three monkeys (M1, M2 and M3; **Figure 1A**). To screen for face-selective neurons, we measured the responses of each cell to a battery of grayscale images of 14 categories (**Figure 1B**). Face-selective neurons, such as the two examples in Figure 1B, were defined as those that showed at least a twofold greater response to faces than bodies and fruits. This selection criterion yielded 102 single units in ML and 71 single units in AL. Face-selective cells in AL had a higher face selectivity index than cells in ML (**Figure 1C**; *Mdn*_ML_ = 0.60, *Mdn*_AL_ = 0.92, Mann-Whitney-Wilcoxon rank-sum test U=1842, *p*<0.001); cells in AL and ML had similar net firing rates (*Mdn*_ML_ = 6.38, *Mdn*_AL_ = 6.28, U=3671, *p*>0.88).

**Figure 1.**
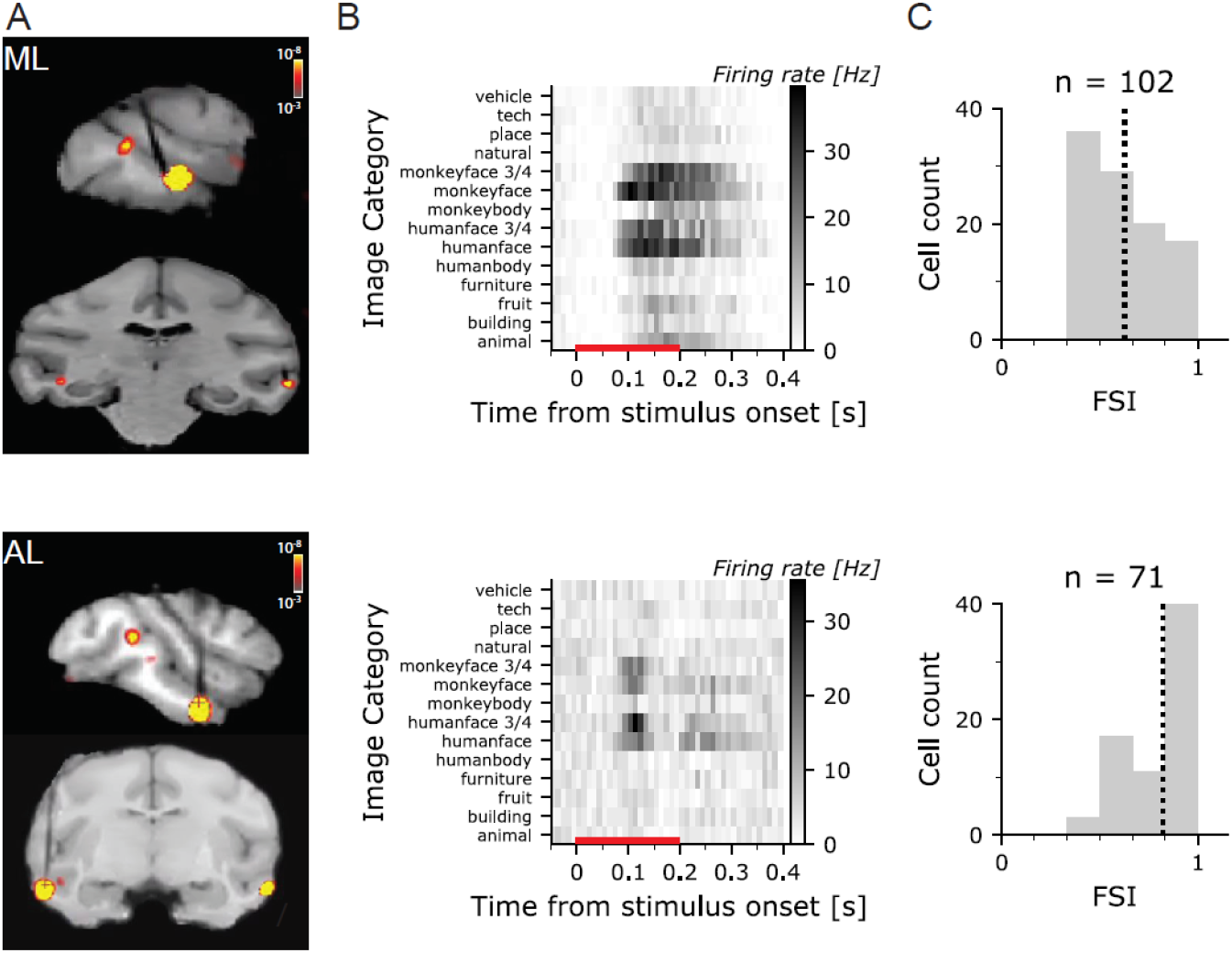
FMRI-guided microelectrode recording of face-selective cells in macaque inferior temporal cortex. **A.** Sagittal MR Images with recording micro-electrodes; superimposed is shown the fMRI contrast maps of faces>bodies uncover the face patches (ML-M2 top; AL-M3 bottom). Electrodes appear black in the MRI images, and target the face patches. **B**. Post-stimulus time histogram for an example cell in ML (top) and AL (bottom) showing the responses to achromatic images used to identify face cells. The red line along the x-axis shows the stimulus duration. **C.** Histogram showing the face-selectivity index (FSI) for the population of face cells in ML (top) and AL (bottom); the dashed vertical line shows the mean FSI. All cells with an FSI>=1/3 were included in the analysis.

We measured responses to color for those cells that showed a face-selectivity index (FSI) greater than or equal to 1/3. Color responses were obtained using monochromatic versions of photographs of faces, bodies, and fruits (**Figure 2A**). Importantly the stimuli were developed with an innovative approach that maintains the luminance contrast of the original images (see Methods). To create a given image in a target color, we replaced each gray value of the original achromatic image with the target hue of the same luminance value. Thus the false-colored images maintained the luminance contrast of the original image. Conventional methods for creating colored images often simply apply a color filter to the image—the procedure replaces the white in the original achromatic picture with a color. The color in images created in that way is easily attributed to the illuminant (or viewing through a colored lens), and not the color of the surface. Moreover, images of different colors created in that way can vary in luminance contrast because of errors in estimating equiluminance, which would confound the interpretation of any color response. Our image-generation methods and stimulus calibration circumvent this potential problem.

**Figure 2.**
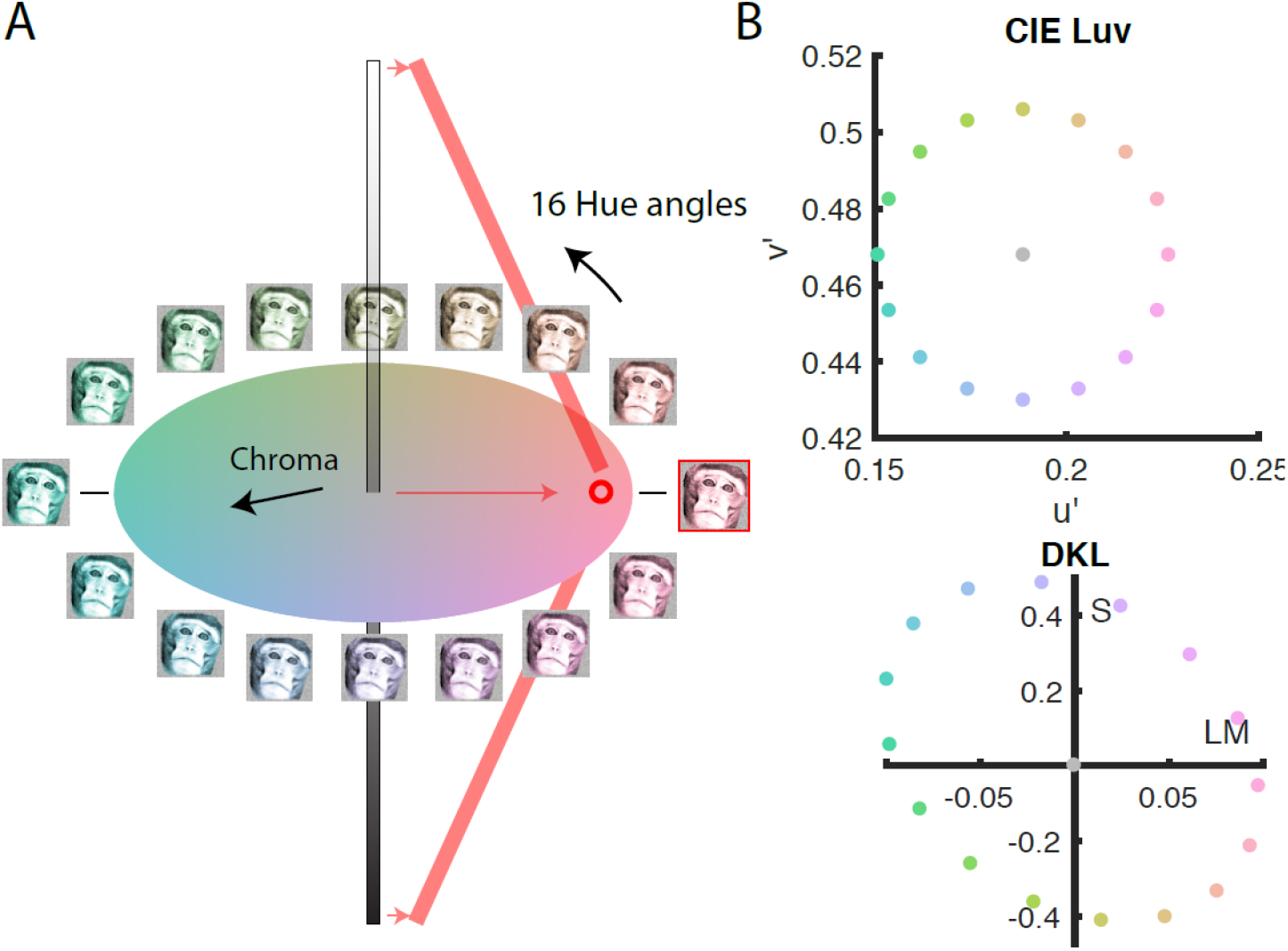
Stimuli used to measure the color responses of face-selective cells in macaque inferior temporal cortex. **A.** Color space illustrating the procedure for generating the colored images. Each colored image was generated by replacing the pixels in the original achromatic image with a given hue that preserved the luminance of original pixel. **B.** Sixteen colored versions were generated; the chromaticity of the most saturated pixel in each version is shown in CIELUV (u*, v*; top panel) and cone-opponent color space (DKL, bottom panel).

To obtain quantitative information about color tuning needed to test for the hypothesized weak color bias in face-selective neurons, and to be able to relate the color responses to comparable data in other brain regions, we needed to measure the color response using stimuli that provide a comprehensive sampling of perceptual color space. To satisfy these requirements, we defined the stimuli in CIELUV color space (**Figure 2B**, top panel); this space captures the representation of color within the V4 Complex (Bohon et al., 2016; Roe et al., 2012) and allows detailed quantitative assessments of color-tuning functions. The chromaticity of the stimuli can be easily transformed into cone-opponent space, which reflects the cone-opponent cardinal mechanisms evident in the lateral geniculate nucleus (Derrington et al., 1984; Sun et al., 2006) (**Figure 2B**, bottom panel).

**Figure 3** shows the responses of a representative sample of six face-selective neurons to the colored images of faces, bodies, and fruit; cells 1-3 were recorded in face patch ML; cells 4-6 were recorded in face patch AL. The top panels show the average responses to images of faces, bodies, and fruits. As predicted given the screening criterion, responses were always substantially larger to faces than to the other stimulus categories, with face-selectivity indices ranging from 0.44 to ^~^1. For each cell we defined a time period for quantification of the responses (blue bar in **Figure 3A, 3B**). We used a single continuous time period for all cells whose duration was tailored to each cell. The response of cells showed complex temporal dynamics. For example, cell #4 showed two peaks (at 105 ms and 225 ms) and the intervening firing rate dipped back to baseline—in cells such as this one, the time period for quantification only included the initial peak. Using multiple time periods for some cells such as cell #4 did not change the main conclusions (data not shown). The median latencies and the duration of the response time periods across the population of cells are reported in **Table 1**. Cells in ML showed a shorter latency than cells in AL within each monkey. But the variability in latencies for cells in ML or AL across monkeys was greater than the difference in latency between ML and AL within any monkey.

**Table 1.** Number of cells, median latency and duration (in ms) of the response window used for the main analysis, by monkey and by face patch.

|  | *M1* |  |  | *M2* |  |  | *M3* |  |  | *Total* |  |  |
| --- | --- | --- | --- | --- | --- | --- | --- | --- | --- | --- | --- | --- |
|  | #cells | latency | duration | #cells | latency | duration | #cells | latency | duration | #cells | latency | duration |
| ML | 45 | 125 | 180 | 57 | 75 | 150 | - | - | - | 102 | 95 | 175 |
| AL | 7 | 135 | 160 | 15 | 95 | 130 | 49 | 75 | 60 | 71 | 85 | 110 |

**Table 2.** Coordinates of the target hues and adaptation point in Cie xyY, CIE LUV and cone contrast spaces.

|  | *CIE xyY* |  |  | *CIE Luv* |  | *DKL* |  |
| --- | --- | --- | --- | --- | --- | --- | --- |
|  | x | y | Y | u' | v' | LM | S |
| white point | 0.3009 | 0.3311 | 56.5 | 0.1889 | 0.4677 | 0 | 0 |
| hue 0° | 0.3473 | 0.3183 | 56.5 | 0.2268 | 0.4677 | 0.0979 | 0.0547 |
| hue 22.5° | 0.3581 | 0.3427 | 56.5 | 0.2239 | 0.4822 | 0.0935 | 0.2145 |
| hue 45° | 0.3607 | 0.3675 | 56.5 | 0.2157 | 0.4945 | 0.0760 | 0.3347 |
| hue 67.5° | 0.3536 | 0.3884 | 56.5 | 0.2034 | 0.5027 | 0.0483 | 0.4026 |
| hue 90° | 0.3370 | 0.4010 | 56.5 | 0.1889 | 0.5056 | 0.0145 | 0.4123 |
| hue 112.5° | 0.3137 | 0.4020 | 56.5 | 0.1744 | 0.5027 | -0.0214 | 0.3637 |
| hue 135° | 0.2882 | 0.3909 | 56.5 | 0.1621 | 0.4945 | -0.0550 | 0.2615 |
| hue 157.5° | 0.2659 | 0.3704 | 56.5 | 0.1538 | 0.4822 | -0.0819 | 0.1164 |
| hue 180° | 0.2505 | 0.3450 | 56.5 | 0.1510 | 0.4677 | -0.0979 | -0.0547 |
| hue 202.5° | 0.2441 | 0.3196 | 56.5 | 0.1538 | 0.4532 | -0.0995 | -0.2282 |
| hue 225° | 0.2465 | 0.2980 | 56.5 | 0.1621 | 0.4409 | -0.0852 | -0.3754 |
| hue 247.5° | 0.2563 | 0.2826 | 56.5 | 0.1744 | 0.4326 | -0.0562 | -0.4678 |
| hue 270° | 0.2717 | 0.2747 | 56.5 | 0.1889 | 0.4298 | -0.0171 | -0.4851 |
| hue 292.5° | 0.2907 | 0.2748 | 56.5 | 0.2034 | 0.4326 | 0.0248 | -0.4226 |
| hue 315° | 0.3111 | 0.2826 | 56.5 | 0.2157 | 0.4409 | 0.0617 | -0.2933 |
| hue 337.5° | 0.3308 | 0.2975 | 56.5 | 0.2239 | 0.4532 | 0.0871 | -0.1239 |

**Figure 3.**
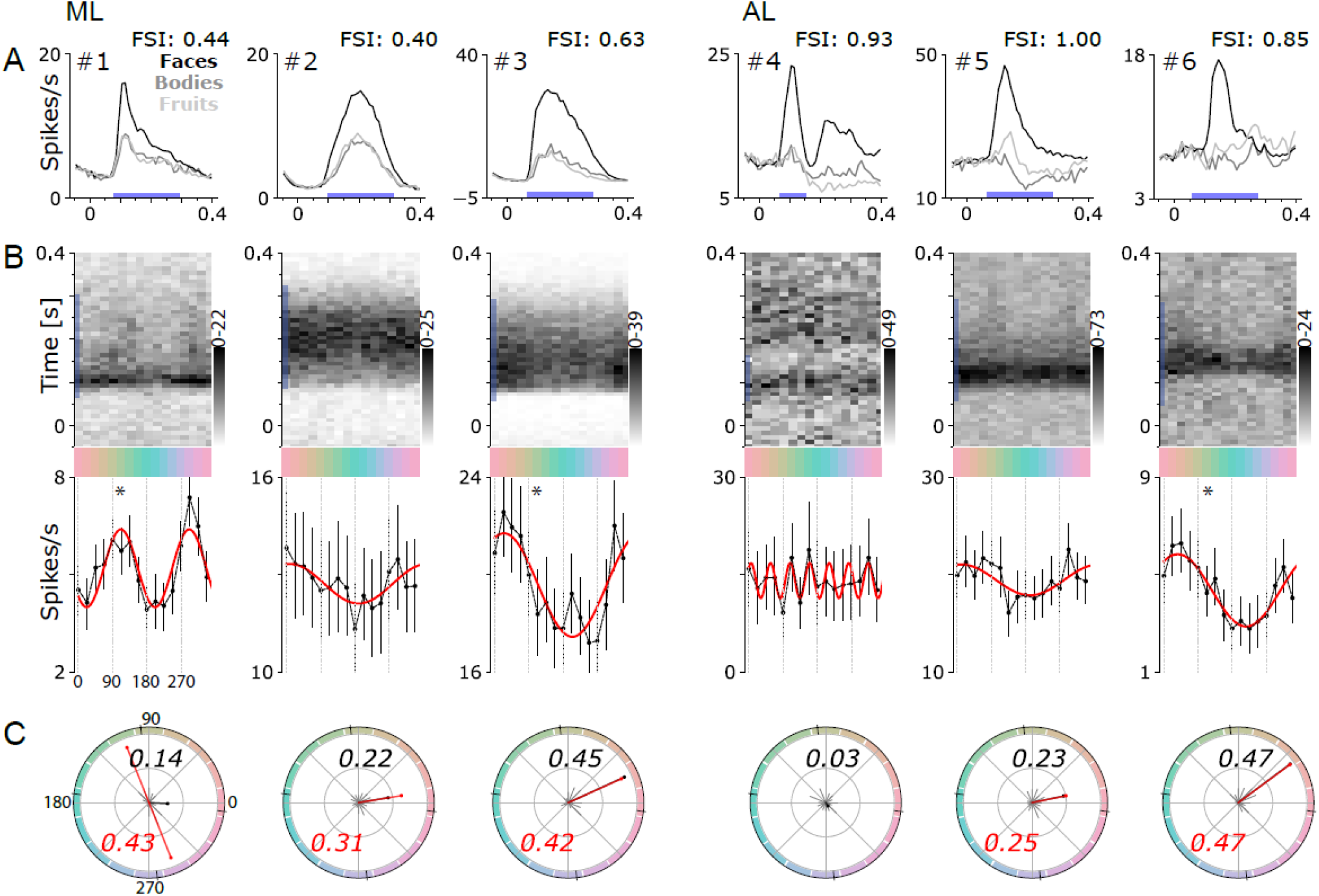
Responses to color of six face-selective cells in macaque inferior temporal cortex (2 cells for each monkey M1: #2, 3 – M2: #1, 5 – M3: #4, 6). **A.** Average responses to images of faces, bodies, and fruits. The time period during which the responses were quantified in subsequent analyses is shown by the blue bar along the x-axis. **B.** Post-stimulus time histogram (top panels) showing the average responses to colored images of faces; blue bar along the time axis as in panel A. Average response to face images of each color (bottom), quantified during the time period indicated by the blue bar in A and B. Error bars show 95% confidence intervals; the red line shows the best fitting sine wave. **C.** Polar plot showing normalized responses to all hues; the sum of the responses to the 16 colors is normalized to equal 1. The bold black text states the norm of the vector sum. The red line shows the normalized amplitude and phase of the best fitting sine wave for neurons whose best fit was a first or second harmonic; the red text states the value of the normalized amplitude of the best-fitting sine. The small black lines on the edges of the circle show the cardinal axes of the cone-opponent color space.

The top panels in **Figure 3B** show post-stimulus-time-histograms (PSTHs) of the responses to the colored faces, averaging across the four different face categories we used (monkey and human faces, frontal faces and ¾-view faces; see Figure 1B). The orientation of the PSTH shows time on the y-axis and image color on the x-axis. The stimulus onset is at 0 seconds, and darker gray corresponds to higher firing rate. The responses of the neurons are delayed by a latency reflecting the time for visual signals to be processed by the retina and relayed through the visual-processing hierarchy to inferior temporal cortex. The cells in **Figure 3** were representative of the population: three of the cells were modulated by the color of the stimulus (cells #1, #3, #6), reflected in the average response over the response window (black trace below the PSTH; see methods for significance calculation). Among the population of face-selective neurons, 25% were significantly modulated by color (23/102 cells in ML; 21/71 cells in AL); moreover, as described below in the population analyses, this metric underestimates the population color response. Cells #2, #4, and #5 were not modulated significantly by color. The red traces in Figure 3B show the best-fitting sine wave following Fourier analysis of the color responses, described below.

Note that the firing rates shown in the bottom panels of **Figure 3B** are averages over the response window and so are lower than the peak firing rates shown in **Figure 3A**. The variance in firing rate caused by changes in color was modest. For example, the firing rate in cell #3 varied between 18 and 22 spikes/second above background, which corresponds to +/- 10% of the average stimulus-driven response. Across the population, the variation in firing rate due to hue was +/-24% of the average stimulus-driven response.

**Figure 3C** plots the cells’ responses as a function of color angle; the vector sum is shown as the bolded black line, and its norm varies between 0 and 1 (0=equal net firing rate for all hues, i.e. no color tuning; values for each cell are shown in black text). The vector sum provides one method to quantify the color tuning, but it would be misleading in situations where neurons have more than one peak. We overcome that limitation by quantifying the results with a Fourier analysis. We determined the best-fitting sine wave of the color-tuning response for each cell (red lines, **Figure 3B**). Most cells (71/173) were best fit by the first harmonic (a single cycle), which shows that these cells have a single preferred color (see cell #3, #6, **Figure 3**); but some cells were best fit by the second harmonic (21/173), indicative of a preference for a color axis in color space, rather than a single color direction (cell #1, **Figure 3**). The amplitude and phase angle of the best fitting harmonic is shown in red in **Figure 3C**. The color preferences assessed by the norm of the vector sum and the normalized amplitude of the first harmonic were correlated (Pearson r=0.93, p<0.001). Among the 44 cells showing significant color modulation, the power of both the first and the second harmonic was higher than the noise level estimated as the power to the 8^th^ harmonic (red lines, **Figure 4**); 37/44 cells showed highest power to the first harmonic; 5/44 showed highest power to the second harmonic; 1/44, to the third; and 1/44 to the fourth.

**Figure 4.**
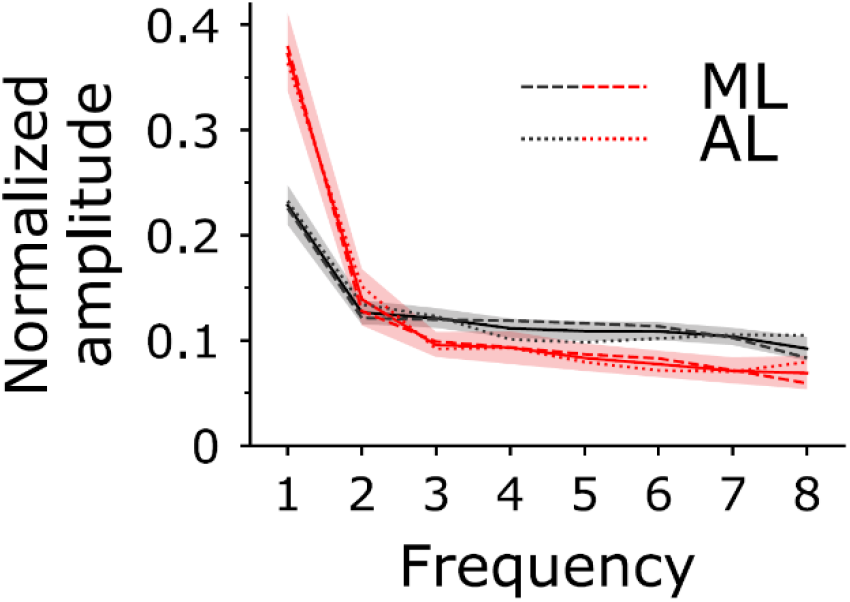
Power spectrum of the Fourier analysis of the color-tuning responses of face-selective neurons in macaque inferior temporal cortex. Average normalized amplitude of each harmonic component for the population (in black) and for significantly color tuned cells (in red). Surrounding shaded areas show 95% confidence intervals and dashed and dotted lines, the averages for respectively ML and AL.

Figures 3 and 4 provide evidence that some face-selective cells were sensitive to color. Among the population, was there a consistent color preference predicted by the hypothesis **Figure 5** shows the color responses of all the face-selective cells rank ordered by the significance of the color tuning, with the most significantly color-tuned cells at the top. Each row shows data from one cell. The gray level shows the normalized response to the given color (the sum of the values across colors for a given cell is 1). Many of the most significantly color tuned neurons in both ML and AL preferred warm colors, as evident by the dark regions on the upper left and right of the panels in Figure 5. But there were some exceptions. For example, cells represented by rows 6 and 7 in the ML panel and rows 1 and 2 in the AL panel of Figure 5 showed a preference for greenish colors.

**Figure 5.**
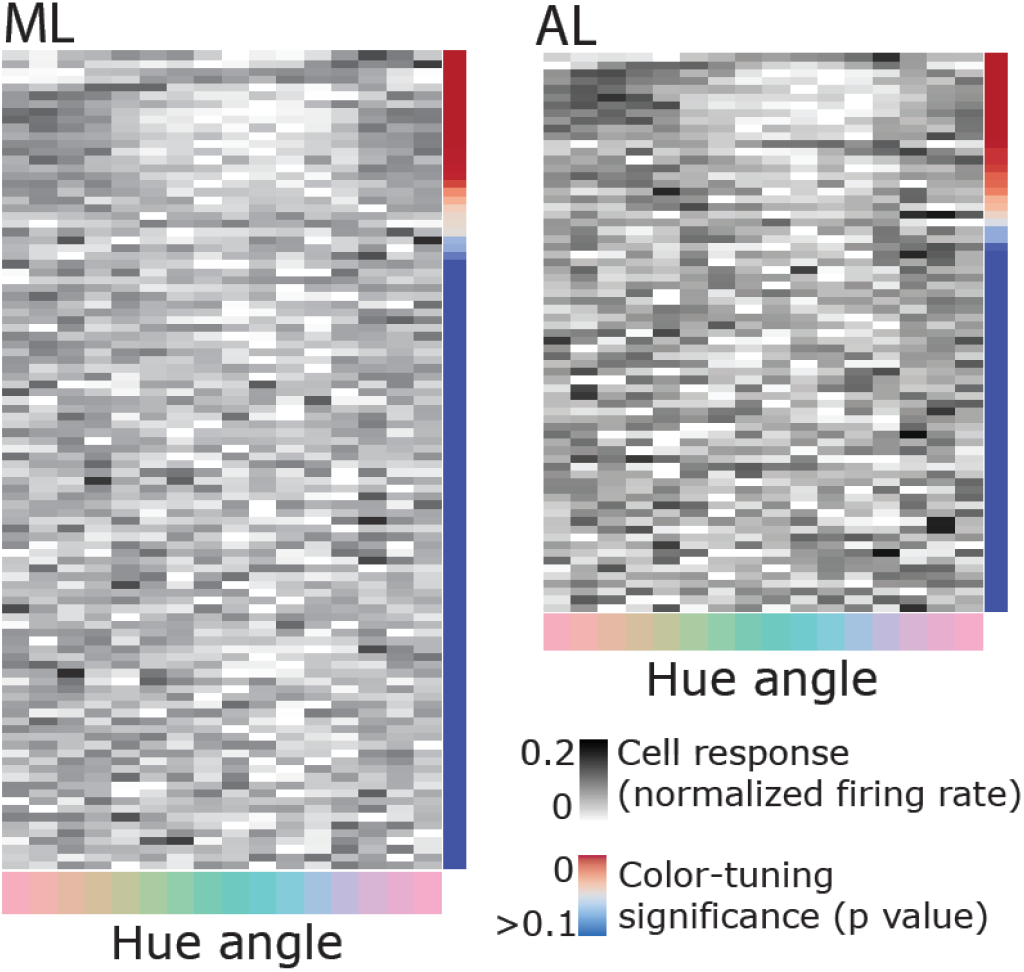
Responses to face images in each of 16 colors, for each cell in the ML face patch (top) and the AL face patch (bottom). Each row shows data for one cell. Cells are ordered from the top by descending color selectivity (p-value indicated by the color scale). The plot shows normalized responses: the sum of the responses to the 16 colors for each row adds up to one (darker gray indicates relatively stronger responses). A maximal preference for one hue would be indicated by black for that hue and white for all others.

**Figure 6** quantifies the color responses of the population of single cells using Fourier analysis. The left panel of Figure 6A shows a polar histogram of the peak color direction for cells with maximum power to the first harmonic; significantly color-tuned cells are shown in dark gray. These results confirm the population bias towards the L>M pole of the L-M cardinal axis, providing the first quantitative evidence in support of the hypothesis that face-selective neurons are color tuned, and specifically tuned to the colors predicted by the face-specific color-shape interaction. This bias is also evident when analyzing the color direction of the best-fitting first harmonic for all cells in the population (including those that did not have maximum power in the first harmonic, **Figure 6B**, left panel). Interestingly, cells with maximum power to the second harmonic showed a phase angle biased for the S-cone cardinal axis (**Figure 6A**, right panel). And this bias was also evident when analyzing the best fitting second harmonic for all cells in the population (including those that did not have maximum power in the second harmonic; **Figure 6B**, right panel).

**Figure 6.**
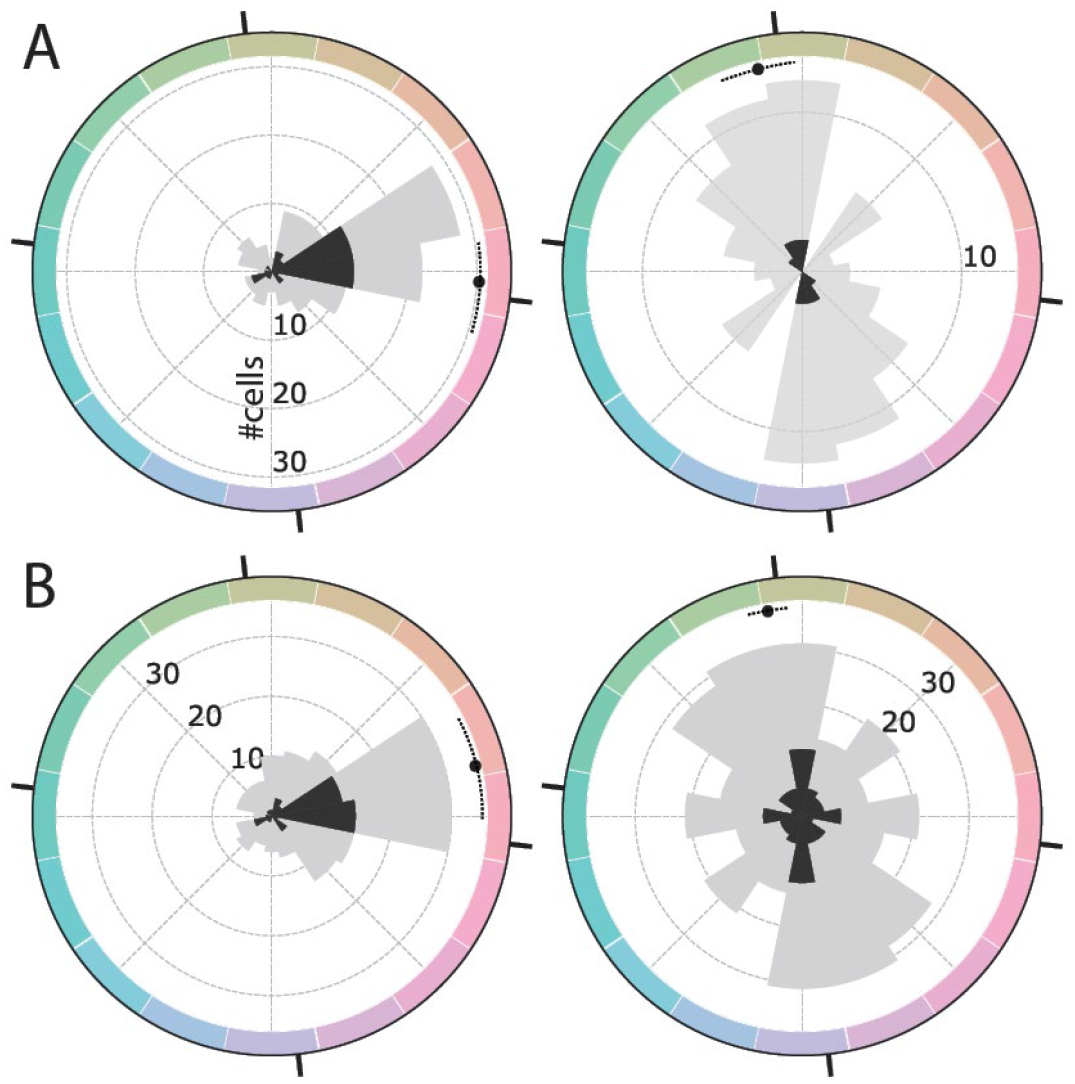
Fourier analysis of the color responses of face-selective cells in macaque inferior temporal cortex. **A.** Left panel shows the distribution of the phase angle for cells with higher amplitude in the first harmonic than the second harmonic (126/173 cells; Mean: −4.40 degrees CI=[−17.0,+8.5]); the right panel shows the phase angle for cells with higher amplitude in the second harmonic compared to the first harmonic (47/173 cells; Mean: +102.7 degrees CI=[+94.5,+111.1]). **B.** Distribution of the phase angle for the whole population (173 cells) for the first harmonic (left panel; mean +13.3 degrees CI=[−0.1,+27.9]) and the second harmonic (right panel; Mean: +99.6 degrees CI=[+94.2,+105.3]). The color space is CIELUV; black tick marks are provided for the cardinal axes of the cone-opponent DKL color space (these are offset from the axes of CIELUV by 6.7 degrees).

How does color tuning relate to color selectivity? If color tuning reflects a computational objective of the circuit one might predict that within the population more color-selective cells will have more consistent color tuning. **Figure 7** quantifies the norm of the vector average (i.e. the peak color preference; y-axis), color selectivity (x-axis), significance of color tuning (symbol gray value), and number of stimulus repeats obtained (symbol size). The data points to the right of the plot converge on 0 degrees (the L>M pole of the L-M axis), consistent with the prediction. Significantly color tuned neurons, defined by the p<0.05 threshold had a mean preferred hue angle that did not differ from insignificantly color tuned neurons (Watson-Williams test, F(1, 171)=0.07, p=0.80) but had a significantly lower variance (**Figure 7**, marginal distribution, Wallraff test, χ^2^=11.59, p<0.001). This effect cannot be attributed to variance in the amount of data collected for different neurons. To prove this, we split the population into two groups, above and below the median number of trials collected per cell. The variance in the peak color tuning was the same for both groups (Wallraff test, χ^2^=1.10, p=0.29).

**Figure 7.**
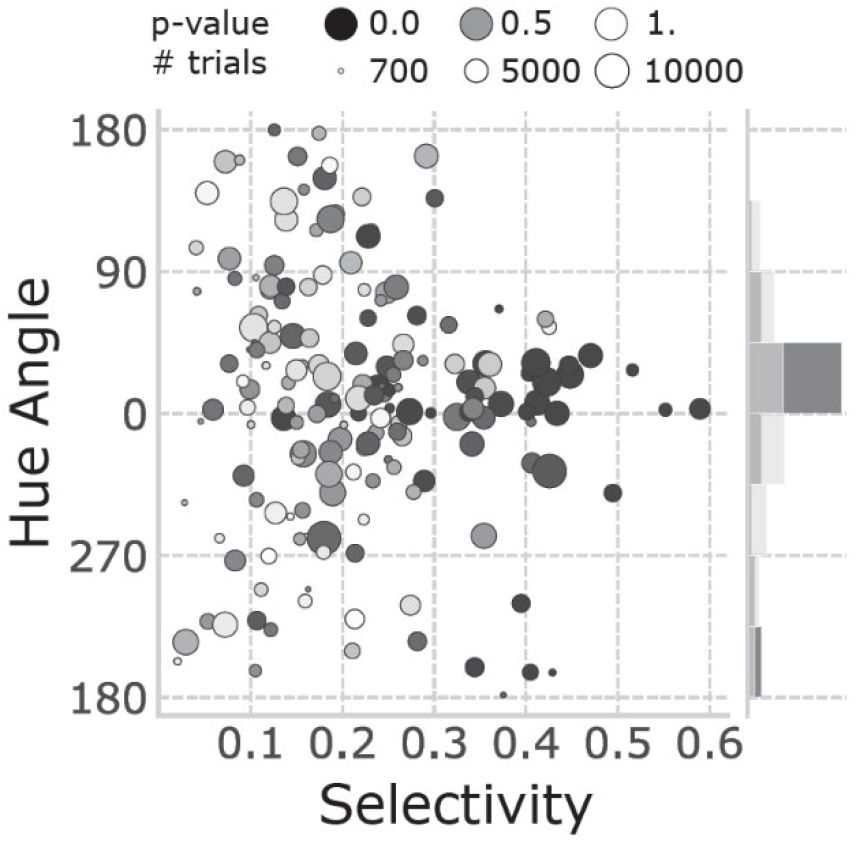
Quantification of the color responses of face-selective cells. Preferred hue angle (direction of the average vector) is plotted as a function of the strength of the color preference (norm of the average vector). Each face-selective cell is represented by a dot. The symbol size corresponds to the number of stimulus presentations. The gray value of the symbols reflects the significance of the color modulation (p-value). The marginal distribution shows the normalized distribution of the preferred hue for significantly (dark grey) and non-significantly (light grey) color-tuned cells.

The face-selective neurons responded strongly to all the colored stimuli, even those of suboptimal color (see Figure 3), which we attribute to the luminance contrast of the stimuli (regardless of the color, the stimuli preserved the luminance contrast of the original images). The luminance-contrast sensitivity of face cells is well-documented (Ohayon et al., 2012). We can directly dissect the role of color and luminance contrast on the cell responses by using equiluminant colored stimuli (**Figure 8A**). These stimuli were created by replacing the range of gray values in the original images with colors of a constant gray value but different saturation: higher luminance gray values were replaced with more saturated color. Responses to these equiluminant stimuli were substantially lower than responses to the colored stimuli that preserved luminance contrast (**Figure 8B**; *Mdn_Iso_* = 0.99, *Mdn_Main_* = 7.64, Wilcoxon rank-sum test U=4643, p<0.001). Nonetheless, the population still showed a response to warm colors (**Figure 8C**). These results show that pure color is not sufficient to strongly drive face-selective cells, but is sufficient to recover the color bias for warm colors predicted.

**Figure 8.**
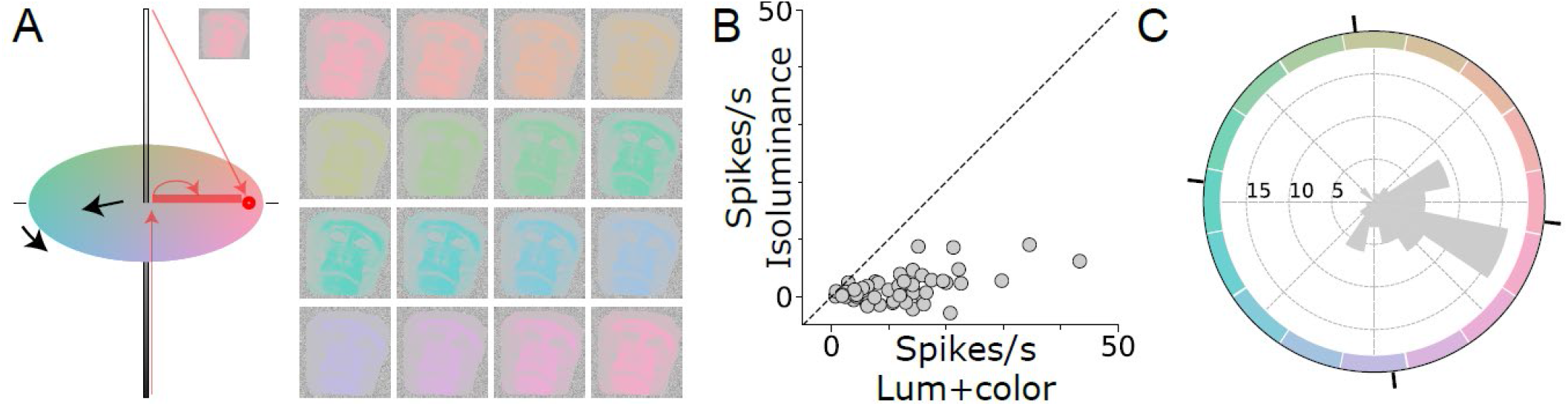
Response to equiluminant stimuli. **A.** Illustration of the construction of an L>M colored image. The range of gray values in the original image were replaced with colors defined by a vector along an equiluminant plane in the color space: white pixels of the original image were rendered in a saturated hue; black pixels were rendered in gray; and gray pixels were rendered in a hue of intermediate saturation. **B.** Firing rate above background of a population of face-selective neurons to equiluminant stimuli (x-axis) versus luminance-preserved colored stimuli (y-axis; N=71 cells), each dot represents one cell. **C.** Histogram of the distribution of peaks in the responses to equiluminant colors. Although the responses to equiluminant colors were very weak, the population nonetheless showed a bias towards L>M colors.

We previously measured the color responses across IT using fMRI (Lafer-Sousa and Conway, 2013; Rosenthal et al., 2018). To directly compare the results of the fMRI with the cell data, we quantified the neural responses within a 400 ms time window starting at the stimulus onset. This time window encompasses the 200 ms duration of the stimulus and the 200 ms inter-stimulus gray period. **Figure 9A** shows the average response for the population, in ML (solid line) and AL (dotted line). The plot shows significant differences among the responses to the colors (non-parametric Friedman test, *χ*^2^= 210.69, p<0.001) and the responses in ML are highly correlated with those in AL (Pearson r=0.89, p<0.001). **Figure 9B** shows the average response across all face-selective cells to face images in each of the 16 colors. This plot underscores two main conclusions: first, responses to all colored images were strong, which we attribute to the fact that all the colored exemplars preserved the luminance contrast of the original achromatic images—the luminance contrast is a main determinant of face-selective responses (Ohayon et al., 2012); and second, among the colored stimuli, responses to the L>M stimuli (appearing pinkish) were higher than response to the M<L stimuli (appearing greenish). The color biases of the population of face-selective cells was strongly correlated with the color biases of the face patches measured with fMRI (**Figure 9C**, Pearson r=0.64 p=0.008).

**Figure 9.**
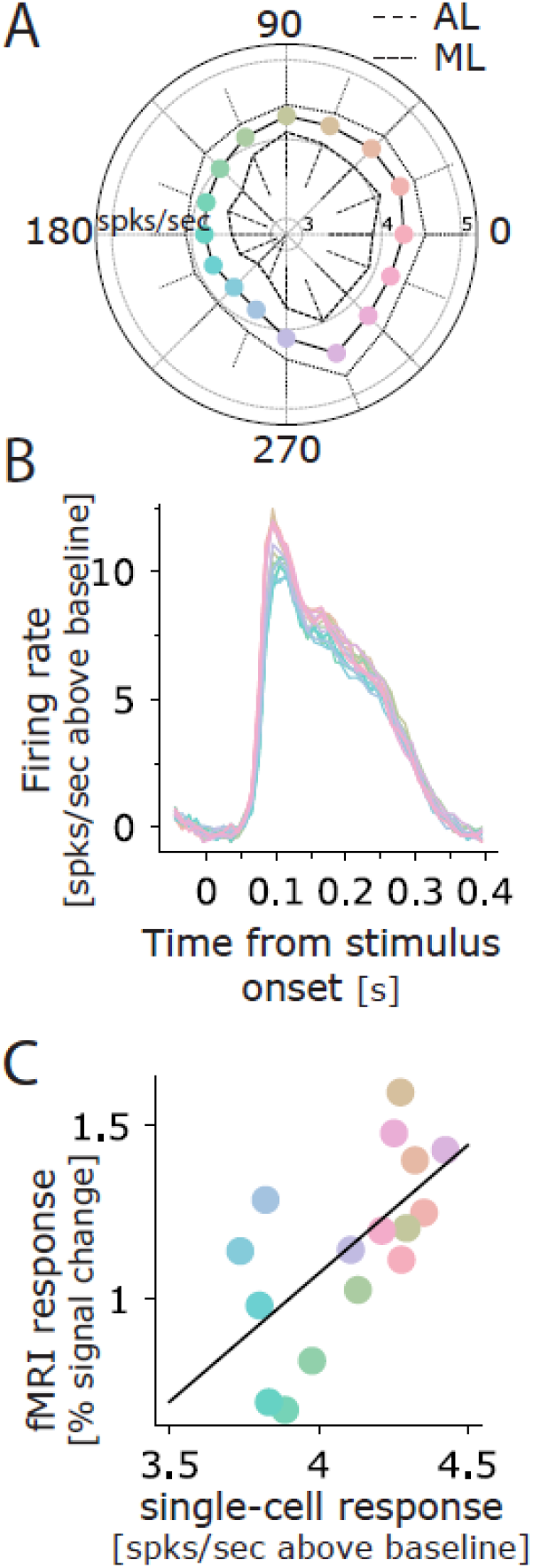
Comparison of color tuning measured using microelectrode recording of single cells in face patches and fMRI. **A.** Average above-background firing rate computed over a 400 ms time window that begins with the stimulus onset. The stimulus duration was 200 ms and there was a 200 ms inter-stimulus interval. **B.** Average above-background response for all face-selective cells (N=173) to face images in 16 colors (the color of the traces corresponds to the colors of the images, see Figure 2). **C.** Correlation between average response across the population of single units and fMRI color tuning assessed in the face patches of monkeys M1 and M2 (see Methods).

**Figure 10.**
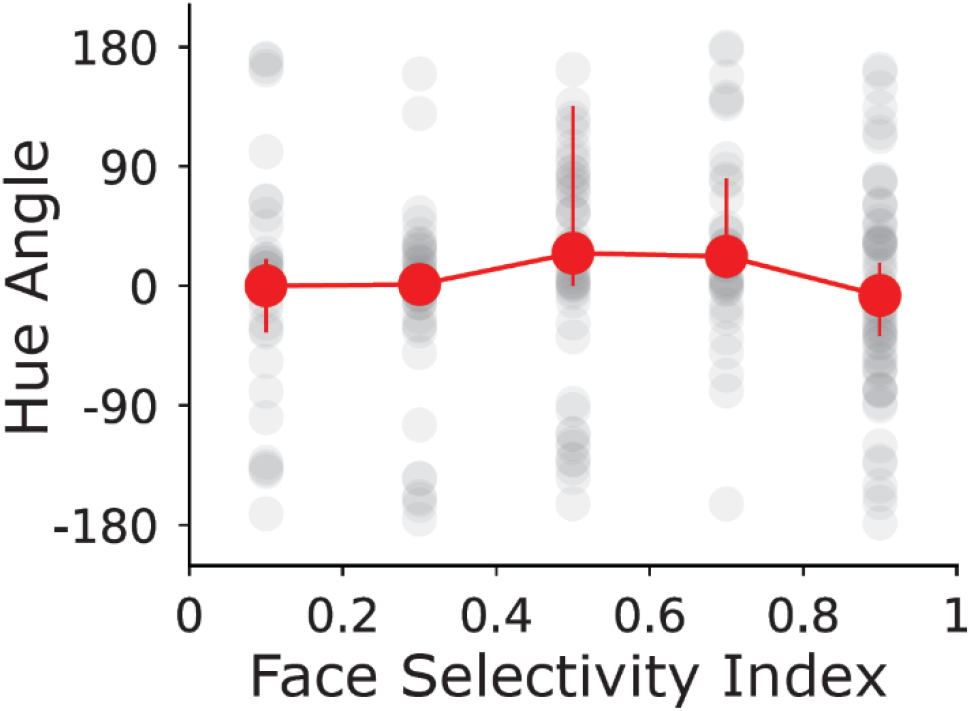
Median hue angles and 95% confidence intervals on the median as a function of Face Selectivity Index are represented in red. Dots represent the values in one bin of 0.2.

## Discussion

Here we document for the first time the color-tuning responses of face-selective neurons in face patches of macaque inferior temporal cortex. The results provide evidence that the population of face-selective neurons shows a bias for warm colors (colors that elicit higher activity in L cones compared to M cones, L>M), supporting the hypothesis we set out to test, namely that face-selective cells encode a prior about the behaviorally relevant component of face color (Hasantash et al., 2019). The color bias was measured using images that preserve the normal luminance contrast relationships of face photographs. In a second series of experiments, we found that face-selective cells were not very responsive to pure color (equiluminant) images of faces, but nonetheless maintained a bias for L>M. The results confirm prior results showing the importance of luminance contrast for face selectivity (Ohayon et al., 2012), and provide single-unit evidence supporting the fMRI observation that face patches respond more strongly to luminance contrast compared to equiluminant color (Lafer-Sousa and Conway, 2013). We speculate that the prior for L>M colors reflects the role of face color as a cue in trichromatic primates, in both humans and macaque monkeys, to sex, dominance, health, and emotion (Conway, 2018; Dubuc et al., 2009; Hiramatsu et al., 2017; Setchell and Wickings, 2005; Stephen et al., 2009; Waitt et al., 2003). The present results uncovering a neural implementation for this prior could provide clues to the neural mechanisms that support these important behaviors. Selective pressures implicating color in social communication have been hypothesized to drive the loss of hair from faces in old-world trichromatic primates (Changizi et al., 2006).

The hypothesis we tested emerged from behavioral data showing a surprising paradoxical effect of memory on face color ((Hasantash et al., 2019), see Introduction). Under conditions that break retinal color vision mechanisms, faces, and only faces, have a paradoxical color appearance: they appear green. That result is analogous to other “anti-Bayesian” phenomena in which a noisy sensory signal gives rise to a perception that is away from the prior, not towards it. These phenomena are perhaps more general than generally appreciated: in addition to face color, they are observed in orientation, biological motion, and size/weight (Brayanov and Smith, 2010; Campbell and Maffei, 1971; Sweeny et al., 2012). As Wei and Stocker showed (2015), the puzzle of these phenomena can be explained by a Bayesian observer model in which image statistics influence both encoding and decoding of sense data (Wei and Stocker, 2015). Wei and Stoker argue that anti-Bayesian phenomena are evidence of neural mechanisms that encode the relevant prior. Hasantash et al show that the same paradoxical face-specific phenomena is evident when participants look at either African American or Caucasian faces, showing that the prior is independent of race. The race-independence further supports the conclusion that the face-color prior is not simply “natural face color” (which can vary along many color directions), but specifically encoded by L>M signals.

The prediction that face-selective neurons should show a bias for L>M has not been directly tested and is not addressable by mining published literature because comprehensive color-tuning functions have not been measured in face-selective neurons. Moreover, the prediction is that the color bias among face cells should be weak because it encodes a prior belief that is only invoked when sense data are ambiguous. Testing this hypothesis therefore requires careful control of the color and luminance statistics of the stimuli. In doing so, we found that 25% of face-selective neurons in both ML and AL had significant color-tuning responses, and the whole population was significantly biased towards L>M colors (**Figure 6**), supporting the hypothesis. The color selectivity of face cells was weaker than the selectivity observed in color-biased domains of posterior inferior temporal cortex (Bohon et al., 2016) and anterior temporal cortex (Komatsu et al., 1992; Namima et al., 2014). As those other studies show, neurons located within color-biased domains identified independently by fMRI typically have narrow (i.e. nonlinear) tuning, and as a population have a complete representation of color space. By contrast, the weak, and biased population color-tuning responses in the face patches is consistent with the notion that the color responses of face cells are not encoding hue generally, but instead a prior about a specific chromatic dimension of face color. It will be interesting to see the extent to which face cells are sensitive to the spatial distribution of color signals within the face; one hypothesis is that manipulating the spatial pattern of color signals across the face to match the perfusion characteristics of skin would enhance the sensitivity of face cells to color. We did not manipulate the spatial aspects of the images in the present experiments to avoid altering the patterns of luminance contrast across the face, patterns to which face cells are very sensitive. It would also be interesting to test the extent to which the cells are sensitive to the distribution of colors across the face, although it is unclear how to do so in a way that allows for a direct comparison with color responses obtained in areas that provide the putative input to the face cells, such as the V4 Complex.

Although most cells had a single peak in the color-tuning function, some face-selective neurons were best fit by two peaks, with maximum power to the second harmonic in the Fourier analysis (**see example cell #1, Figure 3**). The color selectivity of this subset of neurons, assessed as the phase angle of the second harmonic, was aligned with the S-cone axis in color space (**Figure 6**). Is the tuning to the second harmonic meaningful? One might be concerned that these responses reflect a luminance artifact: the estimate of equiluminance may not be accurate especially for colors that modulate S cones, despite appropriate corrections (Vos, 1978). We do not think these responses can be attributed to luminance artifacts because the stimuli preserved the luminance contrast of the original images (the blackest black and whitest white in the colored versions of each image were the same as in the original image). And any residual luminance artifact would be masked by the luminance contrast of the original image.

The color statistics of objects might provide an explanation for the response bias to colors along the S axis. Rosenthal et al. did a comprehensive analysis of the color statistics in a large set of images and showed that animacy can be predicted based only on the extent to which the color of the object modulates S cones (Rosenthal et al., 2018). Thus the tuning to the second harmonic may enable the computation of animacy by IT cells. Regardless of this speculation, to our knowledge, the results provide the first measurements of color tuning biases within extrastriate cortex that reflect the cardinal mechanisms: the first harmonic is tuned along the L-M cardinal axis and the second harmonic is tuned along the S cardinal axis. The cardinal mechanisms correspond to the color tuning of the cone-opponent cells that represent the first post-receptoral stage of color encoding and are reflected in the anatomy and physiology of the lateral geniculate nucleus (Derrington et al., 1984; Martin et al., 2001; Roy et al., 2009; Sun et al., 2006). The cardinal mechanisms are evident in behavioral work that is thought to isolate these subcortical contributions to color vision (Eskew, 2009; Krauskopf et al., 1982). The observation that cortical cells reflect the cardinal mechanisms is surprising because the distinct chromatic signatures associated with the cardinal mechanisms diffuse near the input layers to primary visual cortex (Tailby et al., 2008). The present results show that chromatic signatures corresponding to the cardinal mechanisms reemerge in extrastriate cortical circuits far along the putative visual-processing hierarchy, and they raise the possibility that the behavioral results reflecting the cardinal mechanisms may depend upon cortical, not subcortical, circuits.

Functional MRI response patterns in both macaque monkeys and humans show a multi-stage organizational scheme governed by a repeated eccentricity template, in which color-biased tissue is sandwiched between face-selective tissue (foveal biased) and place-selective tissue (peripheral biased) in parallel streams along the length of the ventral visual pathway (Conway, 2018; Lafer-Sousa and Conway, 2013; Lafer-Sousa et al., 2016). This organization appears to be present early in post-natal development (Arcaro and Livingstone, 2017). The organization might be incorrectly interpreted to imply that the only portions of IT that use color information are the color-biased regions. The results here support the hypothesis that color contributes to operations in most, if not all, of the parallel networks through IT, even those networks that are not explicitly color biased. Establishing in greater detail the relative role of color within these different functionally defined streams in IT is an important objective of future work.

## Methods

### Subjects

Three male rhesus macaques (*Macaca mulatta*), weighing 8-10 kgs, were implanted with an MRI-compatible plastic (Delrin) chamber and headpost. Surgical implantation protocol has been described previously (Lafer-Sousa and Conway, 2013). Designation of the subjects are M1 (monkey 1), M2 (monkey 2), and M3 (monkey 3). Monkeys 1 and 3 (M1, M3) had chambers over the right hemisphere; Monkey 2 (M2) had a chamber on the left hemisphere. All procedures were approved by the Animal Care and Use Committee of the National Eye Institute and complied with the regulations of the National Institutes of Health.

### Functional Imaging targeting of Face Patches

The fMRI procedures we use for localizing face patches have been described (Lafer-Sousa and Conway, 2013; Rosenthal et al., 2018). Two of the animals (M1, M2) are the same as the animals used in Lafer-Sousa and Conway (2013); the face-patch data and color-tuning data are the same as in the earlier reports. Here, we present an analysis of the fMRI color-tuning data restricted to the face patches of inferior temporal cortex. All monkeys were scanned at the Massachusetts General Hospital Martinos Imaging Center in a Siemens 3T Tim Trio scanner. Magnetic resonance images were acquired with a custom-built four-channel magnetic resonance coil system with AC88 gradient insert, which increases the signal-to-noise ratio by allowing very short echo times, providing 1-mm3 spatial resolution and good coverage of the temporal lobe. We used standard echo planar imaging (repetition time = 2 s, 96 × 96 × 50 matrix, 1 mm^3^ voxels; echo time = 13 ms). Monkeys were seated in a sphinx position in a custom-made chair placed inside the bore of the scanner, and they received a juice reward for maintaining fixation on a spot presented at the center of the screen at the end of the bore. An infrared eye tracker (ISCAN) was used to monitor eye movements, and animals were only rewarded by juice for maintaining their gaze within ^~^1 degree of the central fixation target. Magnetic resonance signal contrast was enhanced using a microparticular iron oxide agent – MION (Feraheme, 8–10 mg per kg of body weight, diluted in saline, AMAG Pharmaceuticals), injected intravenously into the femoral vein just before scanning.

Visual stimuli were displayed on a screen subtending 41 by 31 dva, at 49 cm in front of the animal using a JVC-DLA projector (1024 by 768 pixels). The subset of presented stimuli used here to localize face patches consisted in achromatic square photographs of faces and body parts presented centrally on a neutral gray screen (^~^25 cd.m^−2^) and occupying 6 degrees. They were shown in 16 32-s blocks (16 repetition times per block, repetition time = 2 s, 2 images per repetition) presented in one run sequence. The images were matched in average luminance to the neutral gray, maintaining roughly constant average luminance (^~^25 cd.m^−2^) throughout the stimulus sequence. For faces stimuli we used: 16 unique images front-facing of unfamiliar faces (8 human, 8 monkey), The bodies/body parts block comprised 32 unique images of monkey and human bodies (no heads/faces) and body parts. Face patches were localized by contrasting responses to faces versus responses to body parts. 18 runs were used to localize face patches in M1 and M2, and 16 runs were used to localize face patches in M3.

### Physiological recordings

A plastic grid was fitted to the inside of the recording chamber to enable us to reproducibly target regions within the brain, following details reported previously (Conway et al., 2007). We used sharp epoxy-coated tungsten electrodes (FHC), propelled using a hydraulic manual advancer (Narishige). Voltage traces were digitized and saved with a Plexon MAP system (Plexon Inc., Dallas, TX). Spike waveforms were sorted offline with the Plexon Offline Sorter, and single units were defined on the basis of waveform and inter-spike interval.

Recordings were performed in a light-controlled room, with the animals seated in sphinx position. Animals were acclimatized to head restraint in order to minimize head movement during recordings. Animals maintained fixation on a spot on a monitor 57 cm away; the monitor was a CRT Barco subtending 40 by 30 dva, operating at 85 Hz and at a resolution of 1024 by 768 pixels. Eye position was monitored throughout the experiments using and infrared eye-tracker (ISCAN). The monitor was color calibrated using a PR-655 SpectraScan Spectroradiometer (Photo Research Inc.); we achieved 14-bit resolution for each phosphor channel using Bits++ (Cambridge Research Systems).

### Stimuli

Screening stimuli consisted in 10 grayscale exemplars of each of the 14 following categories: animals, buildings, human faces (front view), monkey faces (front view), human faces (3/4 view), monkey faces (3/4 view), fruits, furniture, monkey bodies (no face), human bodies (no face), places, technology objects, indoor places, natural scenes, vehicles. All stimuli were presented on a static luminance noise background of 7.5 by 7.5 degrees of visual angle.

Stimuli for the color-tuning experiments were exemplars of faces of unfamiliar humans and monkeys, front or ¾ view (16 exemplar of frontal view and 8 of ¾ view for each species), fruits known to the monkey (16 exemplars) and bodies/body parts (without head) of unfamiliar humans and monkeys (8 exemplars for each subcategory). For both sets of experiments using colored stimuli (main condition in which the colored stimuli preserved the luminance contrast of the original achromatic images, and the equiluminant condition) we defined 16 target hues equally spaced along the CIELuv color space. For the main condition (Figure 2), each pixel value of the original achromatic image was remapped to the most saturated target color of the same luminance value within the monitor gamut. For the equiluminant condition (Figure 8), each pixel value of the original image was remapped to a pixel on the equiluminant plane, of the same hue as the target but different saturation based on the pixel luminance (see Figure 8).

Note that one way of making false-colored images that has been used in some studies involves the digital equivalent of superimposing a color filter over a black and white picture. In these images, the white of the original image is replaced with a relatively saturated color. Although easy to generate, these false-colored images are luminance compressed compared to the original image: the black in the colored version remains the same luminance as the original, but the white is now a lower luminance than the white in the original. Moreover, it is possible that the estimation of luminance for a given color is inaccurate (see (Conway, 2009) for a discussion). Such inaccuracies would introduce variability in the luminance contrast among the different colored images generated using the color-filter method. The method used presently mitigates the possible impact of chromatic aberration and variability in macular pigmentation, and gives rise to images that are more naturalistic: the images not only retain the luminance contrast of the original images but also appear differently colored, rather than achromatic but viewed through a colored filter (see (Golz and MacLeod, 2002).

### Procedure

One experimental session started by targeting a microelectrode to a face patch, recording a single unit (selected online based on waveform), mapping the receptive field by hand, and choosing the visual-field location that gave the highest response to the screening stimuli. Then we recorded the neural activity while presenting the battery of screening stimuli. If a cell appeared face selective, we then recorded responses to the colored stimuli presented in random order.

### Data Analysis

#### Response window

The response of each neuron was quantified within a response window defined using the average response to all stimuli, in 10 ms bins. The baseline firing rate of each cell was defined as the average response from 50 ms before stimulus onset to 10 ms after stimulus onset. The response window for each cell was determined as one continuous time period initiated when, within two consecutive time bins, the neural response increased above 2.5 standard deviations above the baseline firing rate and terminated either when the response dropped below 2.5 standard deviations of the baseline firing rate in two consecutive bins or after 200 ms following the start of the response window (the shorter of the two options was used). Cells were only included in the analysis if the response window was initiated between 50 ms and 250 ms after stimulus onset, and if the neural response was excitatory (i.e. cells showing suppressive responses to stimulus onset were not included).

#### Face selectivity

The present analysis focused on the color tuning of face-selective neurons. We targeted microelectrode recordings to face patches using fMRI to guide the microelectrode recordings. Face selectivity was assessed using the following index:

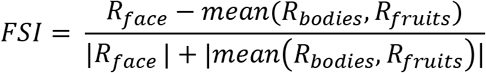

Where R is the average response to stimulus, computed as the difference between the firing rate during the response window and the firing rate during background. FSI values range from −1 to +1, with values above 0 indicating a higher response to faces compared to bodies and fruits. All analyses, with one exception, were restricted to neurons that showed an FSI =>1/3, corresponding to a response to faces at least twice than the response to other non-face stimuli. The exception was the last analysis (see Figure 9), in which we examined the relationship between hue preference and face selectivity. For that last analysis we included an additional 61 cells that were recorded by targeting the same face patches but had an FSI below 1/3.

#### Color Tuning

#### Significance of color modulation

We determined for each cell if there were significant variations in firing rate across the 16 hues by computing the coefficient of variation (the ratio of the standard deviation across hues to the mean) of the data recorded for the neuron compared to the distribution of coefficients of variation obtained by 1000 permutations of hue labels. We considered color modulation to be significant when the p-value was below *α* = 0.05.

#### Vector sum

Color responses were also quantified by determining the vector sum of the color response. This analysis is enabled because hues are circularly distributed: we can consider the neuron’s response to a hue as a vector whose direction is the hue angle, and the vector norm is the strength of the response within the response window compared to baseline. We normalized the vector norms among hues so that the total sum over hues was one. The strength of the hue preference is estimated by the direction and norm of the vector sum. Equation (1) describes the normalized vector computed for each hue; we then sum these vectors using equation (2).

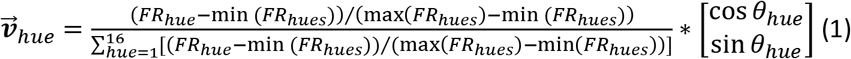

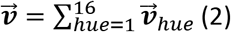

The preferred hue direction of the cell is the angle of 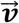. The strength of the hue preference is the norm of 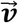, and can take values ranging from 0 (no hue preference) to 1. The norm of the vector sum therefore reflects the narrowness of the color tuning.

#### Fourier analysis

Color responses can be analyzed using Fourier analysis (Krauskopf et al., 1982; Stoughton et al., 2012) that identifies the set of sine waves (frequency, phase angle, and amplitude) that capture the shape of the color-tuning function. We extracted for each cell the normalized amplitude and phase angle of the first 8 harmonics; most of the power was captured by the first two harmonics. The first harmonic has a single peak when plotted in polar coordinates of color space (i.e. a vector pointing to one color); the second harmonic identifies an axis in these coordinates (i.e. the poles of the axis identify an opponent color pair).

#### Correlation between fMRI and electrophysiology

We correlated the single cell activity for each of the 16 hues, as the average firing rate above background over the 400ms corresponding to 200 ms of stimulus followed by 200ms of blank screen for all single cells of all three monkeys, to the average percent signal change to these 16 hues obtained by interpolating from the signal change to the 12 hues used in the fMRI experiment (face-patches were identified over the 2 hemispheres of M1 and M2 of the current study, details of stimuli and ROI definition can be found in Rosenthal et al., 2018).

## Author contributions

The experiments were designed and supervised by BRC, the stimuli were generated by JFD, the neurophysiological data were collected and pre-processed by TG, LT, and SE, the data were analyzed by MD, the paper was written by BRC.

## Acknowledgements

We thank the animal care staff at the National Eye Institute, David Leopold, Chris Baker, Arash Afraz and Rosa Lafer-Sousa for helpful discussion and Isabelle Rosenthal and Theodros Haile for help with the first stages of the experiment. This work was supported by the Intramural research program of the National Eye Institute and the National Institute of Mental Health at the National Institutes of Health.

